# Putamen volume predicts real-time fMRI neurofeedback learning success across paradigms and neurofeedback target regions

**DOI:** 10.1101/2020.10.05.327262

**Authors:** Zhiying Zhao, Shuxia Yao, Jana Zweerings, Xinqi Zhou, Feng Zhou, Huafu Chen, Keith M Kendrick, Klaus Mathiak, Benjamin Becker

**Author notes:** Correspondence to: Klaus Mathiak, Department of Psychiatry, Psychotherapy and Psychosomatics, Medical School, RWTH Aachen University, Pauwelsstr.30, 52074 Aachen, Germany, E-mail address, Benjamin Becker, The Clinical Hospital of Chengdu Brain Science Institute, MOE Key Laboratory for Neuroinformation, High-Field Magnetic Resonance Brain Imaging Key Laboratory of Sichuan Province, University of Electronic Science and Technology of China, Xiyuan Avenue, 2006, 611731 Chengdu, China.

## Abstract

Real-time fMRI guided neurofeedback training has gained increasing interest as a non-invasive brain regulation technique with the potential to normalize functional brain alterations in therapeutic contexts. Individual variations in learning success and treatment response have been observed, yet the neural substrates underlying the learning of self-regulation remain unclear. Against this background, we explored potential brain structural predictors for learning success with pooled data from three real-time fMRI datasets. Our analysis revealed that gray matter volume of the right putamen could predict neurofeedback learning success across the three datasets (n = 66 in total). Importantly, the original studies employed different neurofeedback paradigms during which different brain regions were trained pointing to a general association with learning success independent of specific aspects of the experimental design. Given the role of the putamen in associative learning the finding may reflect an important role of instrumental learning processes and brain structural variations in associated brain regions for successful acquisition of fMRI neurofeedback-guided self-regulation.

## Introduction

Real-time fMRI-based neurofeedback (rt-fMRI NF) is a brain modulation technique that can non-invasively modulate functional brain activity by allowing individuals to gain control over the neural signal (i.e. Blood Oxygenation Level Dependent, BOLD, activation). An increasing number of studies demonstrated that the successful self-regulation of specific regions or networks can induce behavioral changes in domains associated with the trained neural systems in healthy subjects. Based on these promising findings, an increasing number of rt-fMRI NF studies focused on clinical populations. These investigations demonstrated some promising therapeutic effects after single or repeated sessions of training in patients with mental disorders including anxiety disorder (Scheinost, et al., 2013; Zilverstand, et al., 2015), depression (Young, et al., 2017; Young, et al., 2014), attention deficit disorder (Alegria, et al., 2017; Zilverstand, et al., 2017), substance use disorder (Hanlon, et al., 2013; Hartwell, et al., 2016; Kirsch, et al., 2016; Li, et al., 2013) and schizophrenia (Dyck, et al., 2016; Okano, et al., 2020; Orlov, et al., 2018; Zweerings, et al., 2019).

While these studies demonstrate that healthy subjects and patients can learn volitional control over brain activation and initial clinical studies reported beneficial effects on symptom improvement, the precise behavioral and neural mechanisms that underlie the acquisition of rt-fMRI NF-guided self-regulation remain unclear. Within this context recent reviews in the field have proposed key questions that need to be addressed to promote a more conceptually-based account to rt-fMRI NF, such as to clarify which neural substrates support the acquisition of neural self-regulation (Emmert, et al., 2016), and how the regulation success can be translated into changes in specific behavioral domains and ultimately clinical responses (Hampson, 2017). A recent meta-analysis aggregated results reported by 99 rt-fMRI NF studies and found that only 57 out of the 99 studies observed increased regulation in comparison to baseline, and less than half of the studies found overall improvements on the behavioral level (Thibault, et al., 2018). A similar issue has been previously raised for electroencephalogram-based neurofeedback (EEG NF) (Alkoby, et al., 2018). According to this review, 38% of the participants across the 11 EEG NF studies included were not able to gain regulatory control over their brain activity during training. Accordingly, a better understanding of critical aspects that determine regulation success is essential for the progression of the field.

Recent debates on optimizing rt-fMRI NF training efficacy have been mostly focused on methodological aspects such as the number of training sessions, target region selection (Karch, et al., 2015), novel brain-computer interfaces (Lorenzetti, et al., 2018; Mathiak, et al., 2015), instructions and learning strategies (Sitaram, et al., 2017; Stoeckel, et al., 2014) or the specific form of feedback presentation (e.g. intermittent vs. continuous or implicit vs. explicit, see (Emmert, et al., 2017; Stoeckel, et al., 2014)). In addition to optimizing the efficacy of the training *per se* recently emerging frameworks conceptualizing a precision medicine approach for mental disorders (Insel, 2014) proposed that accounting for individual differences in the patients may represent a promising strategy to optimize treatment selection for specific patient populations or on the individual level. Extending this approach to neurofeedback training by determining factors that modulate or predict learning success could help to identify patients with the highest potential to benefit from the training and thus increase the training efficacy regarding symptom improvement. Moreover, the determination of neural predictors may generally help to further determine the complex processes underlying neurofeedback learning.

With the aim to determine the neural basis that underlies neurofeedback acquisition a recent meta-analysis encompassing data from 12 rt-fMRI NF studies revealed a brain network commonly engaged during training. The major nodes of this network included the dorsolateral and ventrolateral areas of the prefrontal cortex (dlPFC and vlPFC), basal ganglia, anterior insula cortex (AIC), anterior cingulate cortex (ACC), thalamus and visual associative areas (Emmert, et al., 2016). A review by Sitaram et al. additionally proposed functional domains and associated brain systems that mediate neurofeedback learning. The authors proposed a network similar to that determined in the aforementioned meta-analysis which supports the involvement of different cognitive systems including executive control, salience detection and reward processing during neurofeedback learning (Sitaram, et al., 2017). In particular, the authors emphasized a key role of the dorsal and ventral striatum because of their central roles in instrumental and associative learning which are highly related to the acquisition of feedback-dependent behavioral modification (Christoffersen and Schachtman, 2016; Gruart, et al., 2015; Yin, et al., 2009).

NF training shares a strong learning-related component with other forms of treatments that aim at modifying behavioral maladaptations, such as cognitive training elements of behavioral therapy. Cognitive-behavioral therapy (CBT) for instance applies learning-based strategies and has been shown efficient to attenuate behavioral and neural dysregulations in a range of mental disorders. Previous studies that combined the personalized medicine approach with neuroimaging in this context have reported that neural recruitment during baseline (Klumpp, et al., 2013) as well as individual variations in brain morphometry – in particular gray matter volumes (Bryant, et al., 2008) in the dysregulated pathways – are predictive for the subsequent treatment response to CBT. When targeting patient populations, rt-fMRI NF training is commonly focused on restoring the control over emotions using mental strategies which share similar components to CBT training (Linhartova, et al., 2019). Results from previous EEG-neurofeedback learning studies suggest that in particular brain structural variations may represent a promising candidate index for predicting learning success. These studies reported for instance that volumes of the cingulate cortex, AIC and putamen as measured by voxel-based morphometry (VBM) were predictive for learning outcomes (Enriquez-Geppert, et al., 2013; Ninaus, et al., 2015).

Given that highly similar cognitive processes may underlie the acquisition of self-regulation by neurofeedback from EEG and fMRI modalities it is thus conceivable that brain structural indices may predict learning success in rt-fMRI NF experiments. A previous meta-analysis of NF studies aimed at determining whether pre-training activation within the target regions of the training collected from more than 400 participants could predict subsequent learning success, yet did not find a common predictor for neurofeedback efficacy, suggesting an examination of more stable alternative predictors of neurofeedback learning success (Haugg, et al., 2020). Against this background, the current study explored whether regional brain volume could predict learning success during rt-fMRI NF training in healthy subjects. Based on the role of the dlPFC, ACC, AIC and striatum in neurofeedback training as suggested by the previous literatures (Emmert, et al., 2016; Sitaram, et al., 2017), we hypothesized that gray matter volumes (GMV) in these regions are associated with neurofeedback learning success in healthy individuals. Furthermore, to increase statistical power and generalize the association across different target regions our hypothesis was tested in data pooled from three datasets targeting different brain regions or pathways with NF.

## Methods

### Datasets and participants

The datasets reported in the current study were collected in healthy samples and were previously published in peer-reviewed journals (Mathiak, et al., 2015; Yao, et al., 2016; Zhao, et al., 2019; Zweerings, et al., 2018) Only data from the experimental neurofeedback runs was included in our analysis (i.e. given the lack of learning success during the sham/control conditions the corresponding data was excluded from the current analyses; see Table 1 for demographics). The datasets are distinct from each other by the experimental design and the targeted training regions.

**Table 1.**
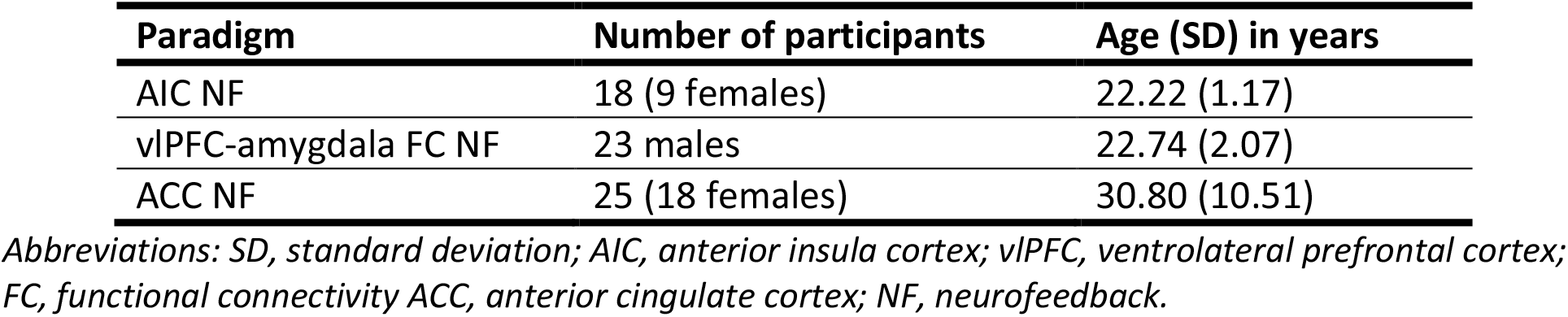
Demographic information of the datasets

Two out of the three datasets were collected on a 3-Tesla MRI system (MR750, General Electric Medical System, Milwaukee, WI, USA) in China. One of them trained up-regulation of AIC activity (Yao, et al., 2016) and the other study trained participants to up-regulate functional connectivity between amygdala and ventrolateral prefrontal cortex (vlPFC) (Zhao, et al., 2019). Both studies found significant change in the trained neural activity after 4 runs of training. T1-weighted brain structure images were collected immediately before the neurofeedback training. The third dataset was collected on a 3-Tesla Siemens MRI system (Magnetom TRIO, Siemens Medical Systems, Erlangen, Germany) at RWTH Aachen University, Germany. Participants in this dataset were collected in two different studies (Mathiak, et al., 2015; Zweerings, et al., 2018) that shared the same training procedure during which participants received 3 sessions of neurofeedback training to increase brain activity in the anatomically defined anterior cingulate cortex (ACC). The imaging parameters and further details of the datasets are provided in the Supporting Information (SI).

### Measurement of neurofeedback learning success

Significant training effects on neural indices on the group-level have been previously demonstrated in all three up-regulation training datasets. **In the current study, we further explored whether variations in brain structure is linked to variations in the subsequent NF learning success on the individual level. Learning success was defined as changes in brain activity in the desired direction (as determined in the original studies) by subtracting the measurements in the early training runs/sessions from the ones in the later stages of the training**. To this end, beta-estimates within the trained regions of interest (ROI) were extracted from the generalized linear models provided by the authors of the original studies. For the Chinese datasets, learning success was previously determined by comparing differences in the targeted brain activity between the first two and last two neurofeedback runs (Yao, et al., 2016; Zhao, et al., 2019). This measurement was directly adopted in the current study. For the ACC regulation data, the learning success was calculated as the difference in mean ACC activation during regulation between the third and the first training sessions, a measure that approximates the approach in the other two datasets (Yao, et al., 2016; Zhao, et al., 2019). The data was extracted with the tailored ACC anatomical mask used during the original neurofeedback training (Mathiak, et al., 2015; Zweerings, et al., 2018). To enable cross-dataset comparisons, the learning success calculated from each dataset was further standardized using the *zscore* function (Rance, et al., 2018) incorporated in MATLAB (R2020a. Natick, Massachusetts: The MathWorks Inc.).

### Preprocessing of the imaging data

The brain structural images were preprocessed with the VBM8 toolbox (http://dbm.neuro.unijena.de/vbm/). Individual brain images were first segmented into gray matter, white matter and cerebrospinal fluid tissue probability maps by unified segmentation as implemented in SPM12 (“New Segment Toolbox”). After segmentation, the gray matter images were non-linearly normalized to the MNI standard space with the DARTEL algorithm (Ashburner, 2007) while accounting for individual whole brain volume sizes (“Modulated normalization” in VBM8) to generate the gray matter volume (GMV) maps. Finally, these gray matter images were smoothed with a 4mm full width at half maximum (FWHM) kernel. Total intracranial volume (TIV) was calculated and subsequently included as a covariate in the regression model. The structural data were initially screened for data quality indices (segmentation, registration and covariance) before being subjected to further analysis.

### Structural predictors for neurofeedback learning

Associations between brain volume and neural learning success were examined by multiple regressions in SPM12 (www.fil.ion.ucl.ac.uk/spm/software/spm12/) using the gray matter images generated in the last step above. In addition to the learning success measurement, dataset, scanner, age, gender and TIV were included in the regression model as control variables.

Our goal was to examine which brain structures proposed in the previous literature might contribute to individual differences in neurofeedback learning. To this end, the correlation between gray matter volume and learning success was separately tested for the core systems of the neurofeedback network, specifically the bilateral dlPFC, ACC, AIC, and striatum (both dorsal and ventral division, details of the masks used for the region of interest, ROI, analysis are provided in SI) with a small volume correction (SVC) approach. Clusters that survived p<0.05 while correcting for family-wise errors (FWE) were considered significant. In order to further describe the determined regions associated with subsequent NF success on the functional network level, we additionally explored the intrinsic functional profile of the cluster that survived the correction. To this end we performed a resting-state functional connectivity (rs-FC) analysis in an independent sample of N = 252 healthy young adults (see SI for more details of the sample and analysis; sample description also in Liu et al., 2020). This analysis employed the identified cluster associated with NF learning success from the VBM analysis as seed region in a seed to whole brain resting state FC analysis.

## Results

### Demographics

After screening for data availability and quality, a total of 66 participants were included in the final analysis. The demographic information of these participants is reported in Table 1. None of them received control/sham feedback during the training.

### Associations between brain structure and learning success

Applying region-wise SVC on the statistical map revealed a significant positive correlation between the volume of a cluster located in the striatum and learning success in the pooled data (t = 4.56, pFWE = 0.039, voxels = 62). No significant associations within the other ROIs emerged. The cluster was located in the dorsal division of the right striatum, primarily in the putamen (x-y-z MNI-coordinates of the peak: 27, 9, 8). No negative correlation between GMV and learning success was observed at the same threshold. Of note, because of the exploratory nature of the current work, no correction for multiple comparisons was performed for the number of tests/masks (N = 4 ROIs tested, see Discussion). The whole-brain map of the structural correlation with learning success is shown in Figure 1, further suggesting an association between learning success and structural variations in the bilateral dorsal striatum at an uncorrected display level (p < 0.001, k > 20).

**Figure 1.**
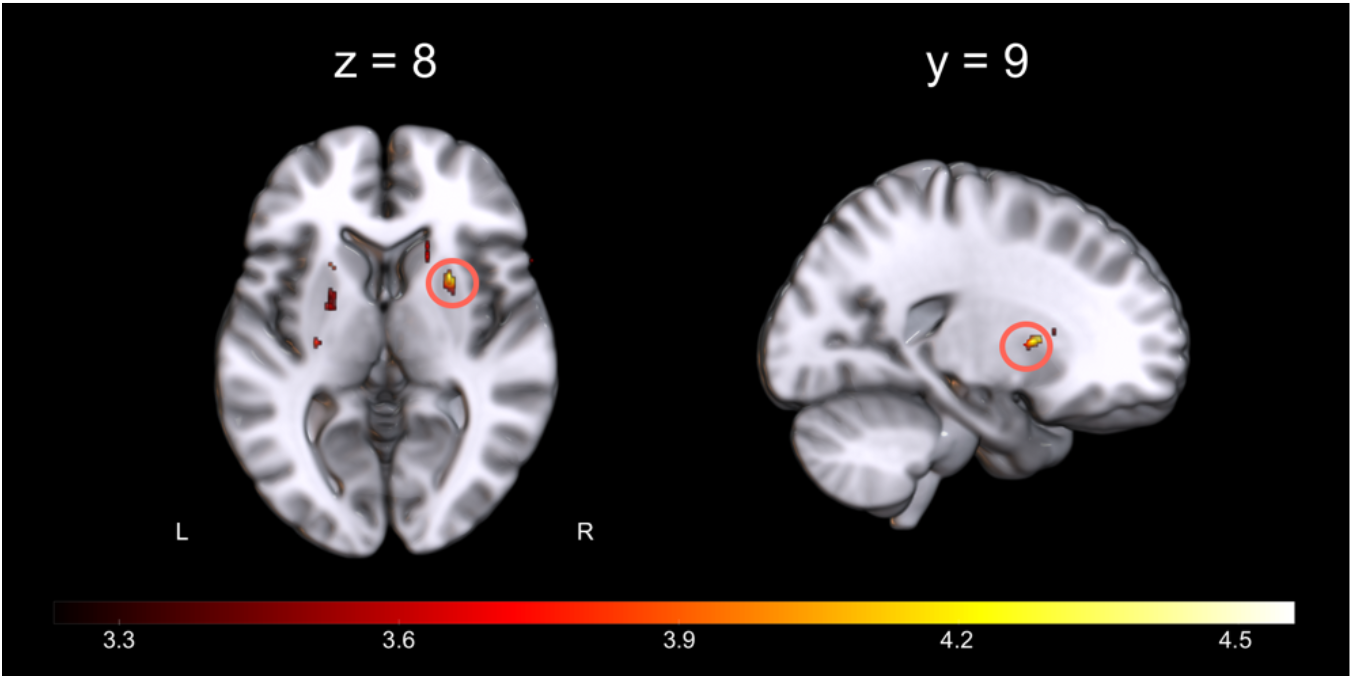
Clusters with a positive association between gray matter volume and neurofeedback learning. Associations between gray matter volume and learning success (displayed uncorrected p < 0.001 with a cluster threshold of k > 20, whole brain level). The circle indicates the putamen region which survived small volume correction in the multiple regression analysis. Other clusters shown in this map are reported in Table S1.

To separately explore the GMV-learning success association pattern in each dataset, regional volume was extracted from the putamen cluster within a 4mm-radius sphere centered at the peak coordinate. Extracted estimates were visualized with a scatter plot (Figure 2). Pearson correlation analysis revealed a consistent pattern of positive associations between larger gray matter volume in this region with better learning success across samples. Interestingly however the GMV estimates varied between the cohorts which may be explained by differences in the different gender ratios between the original studies. Such sex-differences in the morphology of the basal ganglia have been reported previously (Ahsan, et al., 2007). Together our results suggest an association between larger putamen volume with better neurofeedback learning success on the neural level across the studies.

**Figure 2.**
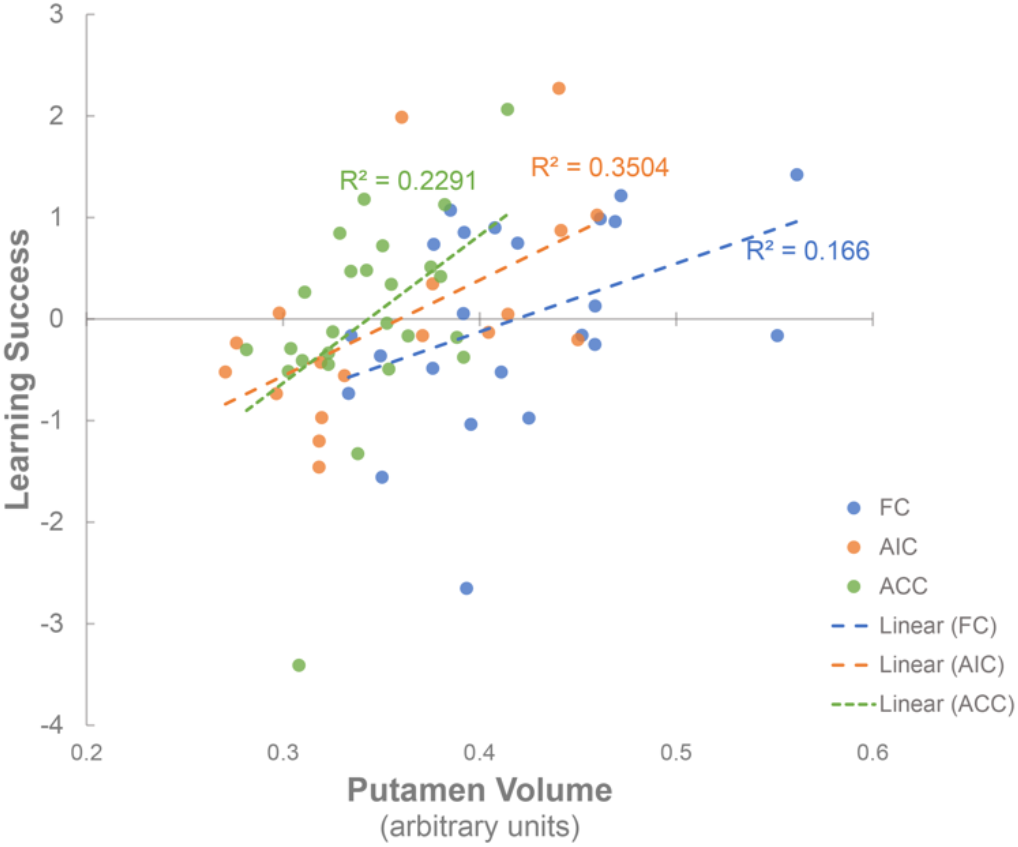
Association between gray matter volume in the putamen area and learning success in the separate datasets. Associations in the separate datasets with data points from each sample coded in different colors. R2 of the linear regressions were denoted. FC = functional connectivity training study, AIC = anterior insula cortex training study, ACC = anterior cingulate cortex training study.

### Network level characterization of the identified region

To facilitate a functional characterization of the identified putamen region we further examined rs-FC of the putamen region (cluster after SVC correction) in a large independent sample (N = 252). The analysis revealed that the identified putamen region exhibited widespread positive intrinsic connectivity with a bilateral network encompassing supplementary motor regions, dlPFC, ACC, AIC, parietal lobe, thalamus, and cerebellum (Figure 3, see also Figure S2 for brain regions that exhibited negative connectivity with the putamen seed region). This network overlapped with the neurofeedback network proposed in the previously literature (Emmert, et al., 2016).

**Figure 3.**
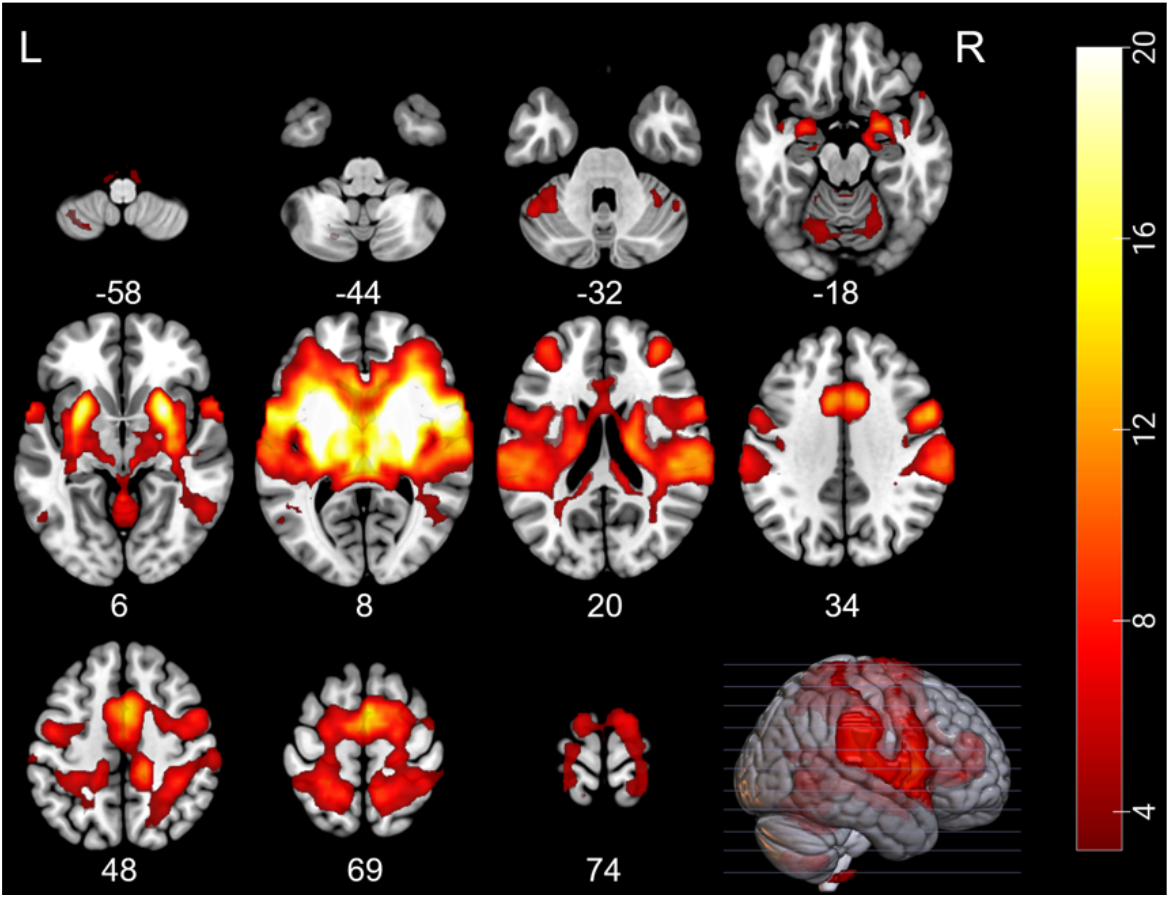
Intrinsic connectivity networks of the identified putamen region. Functional characterization of the identified putamen regions by means of resting state network analysis in an independent sample (N = 252). The displayed map is corrected at cluster level for family-wise error (p_FWE_ < 0.05, cluster forming with height threshold at p < 0.001, k > 68).

## Discussion

In the current study, we investigated whether gray matter volume within the previously proposed neurofeedback training-related brain network could be linked to individual variations in neural regulation success acquired during fMRI-guided neurofeedback training. We found that in three independent samples, better neurofeedback learning success could be predicted by larger volumes of the dorsal striatum, despite differences in experimental designs and target regions employed in the original studies. Moreover, further functional characterization of the identified region by means of determining resting-state functional connectivity between this region and the rest of the brain in an independent sample revealed strong intrinsic functional coupling between the identified dorsal striatum region and other major nodes of the neurofeedback network proposed in the literature (Emmert, et al., 2016; Sitaram, et al., 2017), which further reflects that the dorsal striatum may represent a core node within the networks that support the acquisition of neural self-regulation via fMRI-guided neurofeedback.

The underlying mechanisms and factors that contribute to individual differences in NF learning success have only seldomly been examined. A recent meta-analysis that aimed at identifying functional brain markers has failed to find one that could reliably predict neurofeedback learning success across studies (Haugg, et al., 2020). In the light of these efforts, we asked whether individual variations in brain structure could predict NF learning outcome in three samples and found a positive association between volume of the dorsal striatum, specifically the putamen, and learning success. The findings align with findings from initial studies reporting that learning during rt-fMRI NF has been linked to the recruitment of the striatal system (Mathiak, et al., 2015), probably reflecting a feedback- and reward-related teaching signal (Bray, et al., 2007). A previous study further demonstrated that the failure to acquire fMRI-guided neural self-regulation in patients with schizophrenia was mediated by dysfunctional feedback-related responses in the striatum (Dyck, et al., 2016) and reduced putamen volumes have been repeatedly reported in this population (Gaser, et al., 2004). Additional support for an association between individual variations in brain structure and neurofeedback learning success comes from EEG-neurofeedback studies which may share key mechanisms of the acquisition of neural self-regulation with fMRI-guided NF protocols. Using machine learning approaches a previous EEG-neurofeedback study demonstrated that the fractional anisotropy (FA) value in local fiber structures including cingulum fibers was highly correlated with neurofeedback performance in a sample of twenty healthy participants (Halder, et al., 2013). Two other studies reported that brain volumes of the cingulate area and the putamen along with regions including AIC and prefrontal cortex could predict participants’ self-regulation performance during EEG rhythm training (Enriquez-Geppert, et al., 2013; Ninaus, et al., 2015). Partly resembling these previous findings our results suggest that the structural integrity of the putamen, a brain structure involved in feedback-based and instrumental learning, is associated with neurofeedback learning success during fMRI-guided NF in healthy individuals.

During a typical neurofeedback training setting, subjects gain regulatory control based on feedback that reflects the neural signal of the targeted region or network. The process of gaining control itself may be considered as intrinsically rewarding process. As such increasing regulatory control is followed by the presence of a reinforcing outcome (i.e. the provision of a rewarding feedback signal upon successful regulation efforts in the desired direction) which resembles a form of instrumental learning (Sitaram, et al., 2017). Evidence from animal studies indicate a crucial role of the dorsal striatum in instrumental, associative and procedural learning. This is especially supported by neuromodulatory effects observed on synapses in dorsal striatal regions during both skill learning and instrumental learning processes (Gruart, et al., 2015; Lovinger, 2010). The dorsal striatum encompasses the caudate and putamen, and facilitates the integration of information input from the associative cortices (Haber, 2016). Furthermore, this region is strongly involved in the tracking of action-outcome contingency during associative learning (Brovelli, et al., 2011; Yin and Knowlton, 2006). In humans, focal lesions of the putamen have found to be associated with impaired performance in punishment-based avoidance learning which is in line with a large body of evidence showing the recruitment of this region during encoding of the outcome (Delgado, et al., 2000; O’Doherty, et al., 2004; Palminteri, et al., 2012; Seymour, et al., 2007). In line with recent overarching conceptualizations proposing the importance of instrumental and procedural learning mechanisms in NF learning, responses to NF-guided neural signals reliably engage this region (Emmert, et al., 2016; Shibata, et al., 2019). In a study which trained up-regulation of the right inferior frontal cortex in adolescents with attention deficit disorder, error-monitoring related activity in the putamen region was increased in a stop signal task in the experimental group compared to the active control group. Additionally, this pre-post change had trend-level correlations with neurofeedback learning performance as well as the improvements in symptoms (Criaud, et al., 2020; Lam, et al., 2020). On the brain structural level, VBM studies have found that the gray matter volume of the putamen could be related to both the skill level of piano playing (Granert, et al., 2011) and the performance of neurofeedback learning in sensorimotor rhythm regulation (Ninaus, et al., 2015), suggesting an association with procedural learning success. Taken together, our results may reflect that during neurofeedback learning, individuals with larger putamen volume might benefit from a better ability in adjusting mental strategies based on perceived feedback contingency or a better procedural learning ability which in turn may have promoted increased learning success.

The present data provides the first evidence that individual variations in the morphology of the dorsal striatum may predict NF learning success. The findings are based on three independent datasets with variation in the experimental design and the selected target regions, and hence suggest generalizability of the findings. However, the role of the putamen during neurofeedback training remains speculative due to the lack of direct support from behavioral data. As we have discussed previously, an overarching neurobiological model of neurofeedback learning is yet to be established. In this context, the exact mechanism of action needs to be further examined with tailored experimental designs and validated in larger samples. As a complex cognitive process, neurofeedback learning is presumably underpinned by the involvement of several brain structures or networks in addition to the region discovered in the current study. For instance, dlPFC and ACC are central for sustaining attention to the stimuli and error monitoring during feedback learning (van Duijvenvoorde, et al., 2008). Findings from a recent closed-loop training study indicated a role of the frontostriatal circuitry in learning to regulate high-dimensional brain activity by bridging metacognition (prefrontal) and reinforcement learning (striatum) substrates which further suggested that neurofeedback learning relies on an integrated brain network rather than a single brain structure alone (Cortese, et al., 2020). On the behavioral level, a meta-analysis examining psychological factors for neurofeedback efficacy has highlighted the influence of attention, motivation and mood on training outcome (Kadosh and Staunton, 2019). Therefore it is worth examining whether the volume of the brain structures supporting these functions can be linked to neurofeedback efficacy in a similar way as demonstrated in the current study. Due to the substantial number of brain regions that may contribute to neurofeedback learning, 4 separate ROIs were tested with our small volume correction. However, the multiple-testing issues could not be resolved by simply applying Bonferroni correction on the number of tests as it would also inflate the false negatives rate. Further evidence is needed to allow for a comprehensive interpretation of our findings.

## Conclusions

To the best of our knowledge, this is the first investigation showing that brain structure factor is predictive for rt-fMRI neurofeedback efficacy. The association between putamen volume and learning success may reflect the key role of instrumental learning processes during neurofeedback training. These findings await examination in further studies providing more comprehensive evidence.

## Supporting information

Supplemental methods and results

## Acknowledgments

This work was supported by the National Key Research and Development Program of China (2018YFA0701400) and the National Natural Science Foundation of China grants (NSFC grant number 31700998 and U1808204), the German Research Foundation (DFG; IRTG 2150), and the German Ministry for Education and Research (BMBF; APIC: 01EE1405A-C). We thank Patrick Eisner for his help with data transferring.

The authors declare no conflict of interest.

