## Supplemental methods and results for "Putamen volume predicts real-time fMRI neurofeedback learning success across paradigms and neurofeedback target regions"

**Regions of Interest**

To test for brain structural predictors of learning success, four different bilateral masks were separately applied for small volume correction. For the dorsolateral prefrontal cortex (dlPFC), a mask was created by means of the neurosynth database for meta-analytic approaches in neuroimaging (neurosynth.org). The meta-analysis was conducted in the Neurosynth database by employing the term 'dlpfc' and applying a height threshold of $z > 3$ to the resulting mask. The anatomical mask of the anterior cingulate cortex (ACC) was based on the region of interest selected for the neurofeedback training (Mathiak, et al., 2015; Zweerings, et al., 2018)which corresponds to the anterior-inferior section of the ACC (circumscribed by $y > 0$ and $z < 0$ in MNI space) from the Automated Anatomic Labelling (AAL) atlas (Tzourio-Mazoyer, et al., 2002). The masks for the anterior insula cortex (AIC) and striatum were derived from the the Human Brainnetome Atlas (Fan, et al., 2016), with the AIC consisting of region 167, 168, 173 and 174 of the atlas, and the striatum consisting of all the basal ganglia subregions except globus pallidus. The striatum was further divided into dorsal and ventral sections based on the literature (Di Martino, et al., 2008; Zhao, et al., 2019) for the purpose of results interpretation and illustration. These structures are illustrated in Figure S1 below.

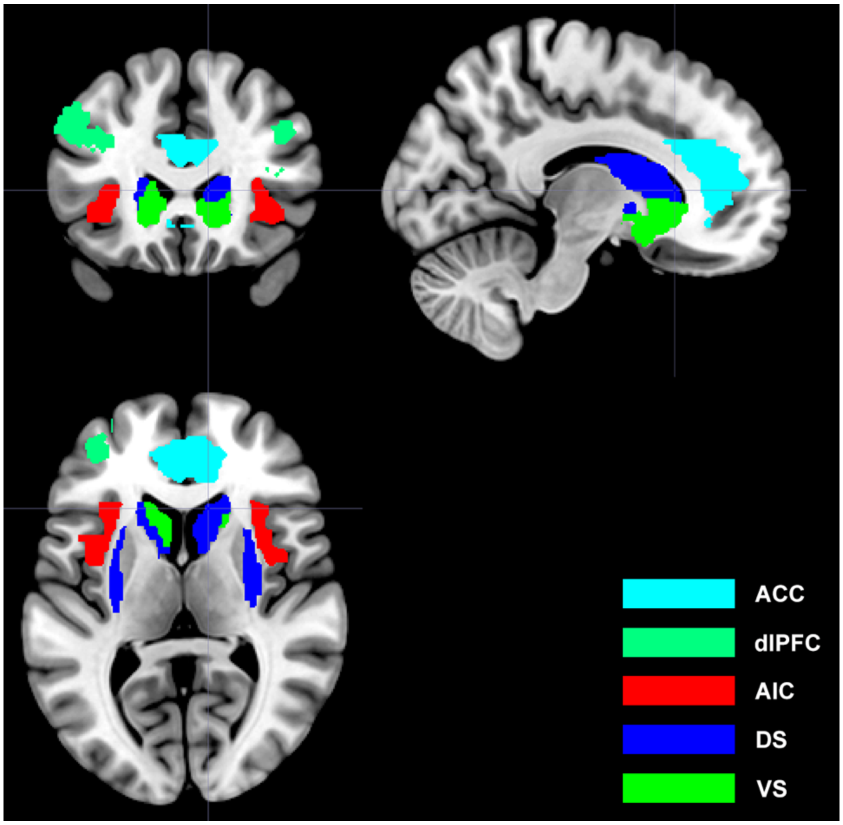

Figure S1. The four regions of interest that were used in small volume correction overlaid on a MNI brain in different colors (see legends at bottom right corner). The striatum is further broken down into ventral and dorsal sections for illustration purpose.
*Abbreviations: ACC, anterior cingulate cortex; dlPFC, dorsolateral prefrontal cortex; AIC, anterior insula cortex; DS, dorsal striatum; VS, ventral striatum.*

**Imaging Parameters**

T1-weighted images in the two Chinese datasets (AIC and FC training) were collected with a FSPGR using the following parameters: TR = 5.97 ms, TE = 1.97 ms, FOV = 256 × 256 mm, slice thickness = 1 mm, flip angle = 9°, 128 slices per volume in sagittal direction. T1 images in the German dataset were collected with the following parameters: FOV = 256 × 256 mm, 176 sagittal slices, 1 $mm^3$ isotropic voxels, TR = 1900 ms, TE = 2.52 ms, flip angle = 8°.

Table S1. Brain regions showed positive correlation with learning success at uncorrected threshold

| Region | Coordinate (x-y-z) | t value | Number of voxels |
|---|---|---|---|
| Globus pallidus | -21, 14, 6 | 4.28 | 23 |
| Dorsal caudate | 15, 16, 20 | 4.06 | 75 |
| Dorsolateral putamen | -28, -18, 4 | 3.95 | 39 |
| Globus pallidus | -22 -3, 10 | 3.80 | 102 |
| Inferior frontal gyrus (BA 44, 45) | 60 16 12 | 3.60 | 20 |

Results thresholded at p< 0.001 (uncorrected), k > 20. *Abbreviation: BA: Brodmann area.*

**Resting-State Data and Analysis**

The resting-state data for computing seed-based functional connectivity with the putamen as seed region were collected in an independent sample of 252 healthy college students in Chengdu, China on a GE 3-Tesla MRI system with a T2* weighted Echo Planar Imaging sequence (TR = 2000 ms, TE = 30 ms, FOV = 240 × 240 mm, flip angle = 90°, image matrix = 64 × 64, 39 interleaved slices with thickness/gap of 3.4/0.6 mm). Data from the same sample have been reported in a previous study (Liu, et al., 2020).

The data were preprocessed and analyzed using SPM12 and DPABI (Data Processing & Analysis for Brain Imaging toolbox, http://rfmri.org/dpabi). The data were corrected for slice timing and head movement and normalized to MNI space with physiological noises from cerebrospinal fluid and white matter as well as global signal removed from the data. Finally, band-pass filtering (0.01-0.08 Hz) were applied to remove unwanted drifts and spikes in the data. Subjects with head movements larger than 2 mm or 2 degrees were excluded from the analysis. After preprocessing, Pearson's correlation was calculated between the timecourses of the sphere (radius = 5 mm) centered at the peak of the putamen cluster (MNI coordinates: 27, 9, 8) and voxels from the rest of the brain. Resulting connectivity maps were entered into a one-sample t-test in SPM12 after Fisher's r-to-z transformation. Negative connectivity map in group-level is presented below.

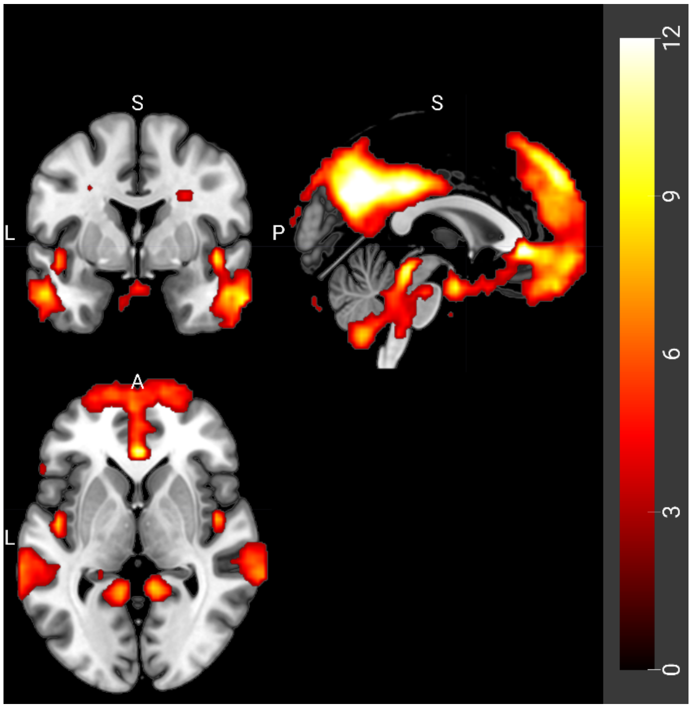


Figure S2. Brain regions showed negative connectivity with the putamen region that predicted learning success. The statistical map is corrected for family-wise errors at cluster level ($p_{FWE} < 0.05$, $k > 68$ at height threshold of $p < 0.001$)
